# Structural determinants of protein kinase A essential for CFTR channel activation

**DOI:** 10.1101/2024.05.27.596024

**Authors:** Csaba Mihályi, Iordan Iordanov, András Szöllősi, László Csanády

**Affiliations:** Department of Biochemistry, Semmelweis University, Tuzolto u. 37-47, Budapest, H-1094, Hungary; HCEMM-SE Molecular Channelopathies Research Group, Semmelweis University, Tuzolto u. 37-47, Budapest, H-1094, Hungary; HUN-REN-SE Ion Channel Research Group, Semmelweis University, Tuzolto u. 37-47, Budapest, H-1094, Hungary

**Keywords:** cystic fibrosis, cAMP-dependent protein kinase, PKI peptide, N-myristoylation, IpaJ

## Abstract

CFTR, the anion channel mutated in cystic fibrosis (CF) patients, is activated by the catalytic subunit of protein kinase A (PKA-C). PKA-C activates CFTR both reversibly, through binding, and irreversibly, through phosphorylation of multiple serines in CFTR’s regulatory (R) domain. Here we identify key molecular determinants of the CFTR/PKA-C interaction essential for these processes. By comparing CFTR current activation in the presence of ATP or an ATP analog unsuitable for phosphotransfer, as well as pseudosubstrate peptides of various lengths, we identify two distinct specific regions of the PKA-C surface which interact with CFTR to cause reversible and irreversible CFTR stimulation, respectively. Whereas the “substrate site” mediates CFTR phosphorylation, a distinct hydrophobic patch (the “docking site”) is responsible for reversible CFTR activation, achieved by stabilizing the R domain in a “released” conformation permissive to channel gating. Furthermore, by comparing PKA-C variants with different posttranslational modification patterns we find that direct membrane tethering of the kinase through its N-terminal myristoyl group is an unappreciated fundamental requirement for CFTR activation: PKA-C demyristoylation abolishes reversible, and profoundly slows irreversible, CFTR stimulation. For the F508del CFTR mutant, present in ∼90% of CF patients, maximal activation by de-myristoylated PKA-C is reduced by ∼10-fold compared to that by myristoylated PKA-C. Finally, in bacterial genera that contain common CF pathogens we identify virulence factors that demyristoylate PKA-C *in vitro*, raising the possibility that during recurrent bacterial infections in CF patients PKA-C demyristoylation may contribute to the exacerbation of lung disease.

**Significance Statement:** CFTR is an anion channel crucial for salt-water transport across epithelia, and is activated by the catalytic subunit of protein kinase A (PKA-C). Reduced activity of mutant CFTR causes cystic fibrosis and CFTR hyperstimulation by sustained PKA-C activity causes diarrhea. PKA-C activates CFTR reversibly through simple binding, and irreversibly by phosphorylating the channel. We uncover here important structural requirements for these two processes. First, two distinct PKA-C surface areas mediate reversible and irreversible CFTR activation. Second, membrane anchoring of PKA-C through a covalently linked fatty (myristic) acid is required for both effects. Finally, we identify bacterial enzymes that cleave the myristic acid from PKA-C, thereby reducing activation of mutant CFTR channels, present in cystic fibrosis patients, by up to tenfold.

## Introduction

The devastating disease cystic fibrosis (CF) is caused by mutations of CFTR, an anion channel expressed in the epithelia of the lung, the pancreas, and the gut, and regulated through phosphorylation by cAMP-dependent protein kinase (protein kinase A, PKA). Defects in the regulation of CFTR by PKA contribute to multiple disease pathologies. On the one hand, profoundly impaired phosphorylation by PKA exacerbates the consequences of protein processing/channel gating defects caused by the most common CF-associated CFTR mutations ΔF508 (1) and G551D (2). Moreover, the potentiating effect of the only FDA-approved CFTR potentiator drug VX-770 is entirely dependent on CFTR channel phosphorylation (3). On the other hand, CFTR hyperstimulation by sustained PKA activity in the intestinal or renal epithelium, respectively, underlies excessive intestinal fluid loss in cholera and other secretory diarrheas (4), as well as progressive renal cyst growth in autosomal dominant polycystic kidney disease (ADPKD) (5). Clinical trials targeting the PKA pathway proved beneficial for the treatment of ADPKD (6).

CFTR is an ATP Binding Cassette (ABC) protein with an anion-selective pore formed by two transmembrane domains (TMD1, 2; Fig. 1A, *gray*) and gated by an ATP hydrolysis cycle at two cytosolic nucleotide binding domains (NBD1, 2; Fig. 1A, *blue* and *green*) (7). Formation of a tight NBD1-NBD2 dimer upon ATP binding opens the pore (8), and disruption of the tight dimer following ATP hydrolysis closes it (9). The two ABC-typical halves of CFTR are linked by a unique cytosolic regulatory (R) domain (Fig. 1A, *red ribbon*) which inhibits channel opening unless it is phosphorylated at multiple serines by PKA (10, 11).

**Figure 1.**
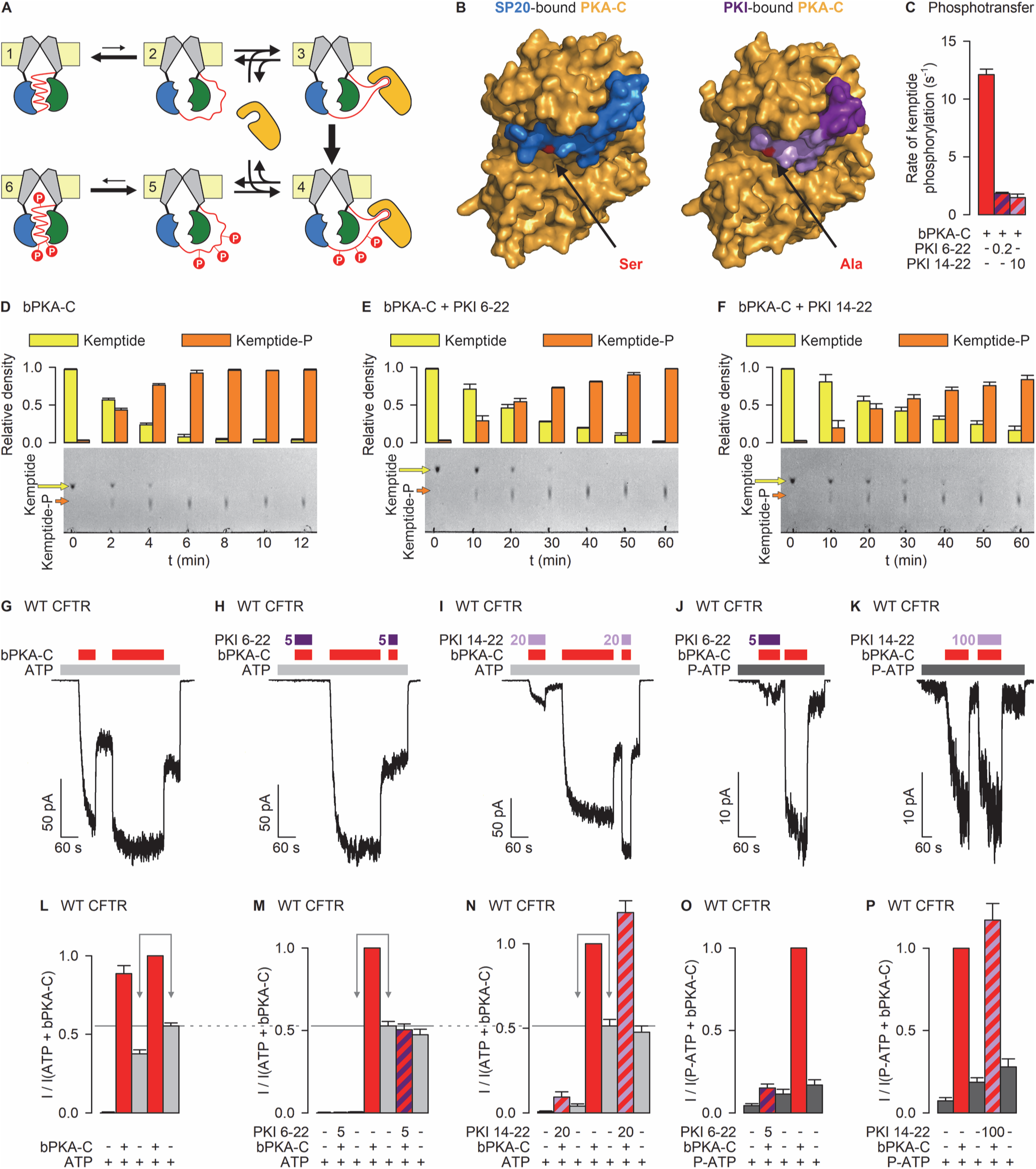
Distinct regions of the PKA-C substrate binding cleft mediate reversible and irreversible CFTR activation. *A*, Cartoon representation of CFTR channel activation. NBD1, *blue*; NBD2, *green*; TMD1 and TMD2, *gray*; R domain, *red ribbon*; phosphate groups, *red circles, “P”*; membrane, *yellow*; PKA-C, *orange*. States 1-6 represent only the possible states of the R domain: in states 2-5 ATP-binding may induce NBD dimerization coupled to pore opening (not depicted). *B*, Surface representations of PKA-C crystal structures with bound (*Left*, PDBID: 4O22) peptide substrate (SP20) or (*Right*, PDBID: 2GFC) inhibitor (PKI(5–24), only residues 6-22 shown). PKA-C, *orange*; SP20, *blue*; PKI residues 6-13, *dark purple*, PKI residues 14-22, *light purple*; target serine in SP20 and corresponding alanine in PKI, *red*. *C*, Estimated *k*_cat_ (s^-1^) for bPKA-C without PKI peptides, with 0.2 μM PKI(6–22)amide, or with 10 μM PKI(14–22)amide, calculated from the data in *D*-*F*. *D*-*F*, (*Bottom*) Time courses of kemptide phosphorylation resolved on TLC sheets. 5 nM bPKA-C without PKI (*D*), with 0.2 μM PKI(6–22)amide (*E*), or with 10 μM PKI(14–22)amide (*F*) were incubated with 20 μM TAMRA-kemptide + 200 μM MgATP for the indicated amounts of time at room temperature. *Upper row of spots*, dephospho-peptide; *lower row of spots*, phospho-peptide. (*Top*) Densitometric analysis of the TLC sheets. *Yellow* and *orange columns* plot fractional densities of the spots corresponding to the dephospho- and phospho-peptide, respectively. Data in *C*-*F* represent mean±SEM, n=3. *G-K*, Macroscopic currents through inside-out patches excised from *Xenopus laevis* oocytes injected with cRNA for WT CFTR. Channels are activated by repeated exposures to 300 nM bPKA-C (*red bars*) in the presence of either 2 mM ATP (*G*-*I*; *light gray bars*) or 10 μM P-ATP (*J*-*K*; *dark gray bars*), with or without PKI(6–22)amide (*H*, *J*; *dark purple bars*) or PKI(14–22)amide (*I*, *K*; *light purple bars*). Colored numbers indicate PKI concentrations in μM. Membrane voltage was −40 mV. *L-P*, Steady-state currents in the sequential segments of the experiment in the panel above, normalized to that in the presence of ATP+bPKA-C or P-ATP+bPKA-C (*red column* (*L*; *2nd red column*)). In *L*-*N* the ratio of the two *gray columns* marked by the *double arrows* report fractional CFTR phosphorylation after 1 minute in 300 nM bPKA-C without PKI peptides (*L*), with 5 μM PKI(6–22)amide (*M*), or with 20 μM PKI(14–22)amide (*N*). Data in *L*-*P* represent mean±SEM, n=7-11 (*L*-*O*), n=4 (*P*).

Because in living cells CFTR’s NBDs are continuously exposed to a saturating ATP concentration, physiological regulation of CFTR activity is realized through R-domain phosphorylation which tightly links the rate of transepithelial anion transport to cytosolic cAMP levels. The PKA holoenzyme consists of two regulatory (R) and two catalytic (C) subunits. Binding to an R subunit keeps the C subunit inactive. When cellular cAMP levels rise, the R subunits bind cAMP and release the C subunits (PKA-C) in free active form, allowing them to bind and phosphorylate their targets (12). PKA-C is a small (∼350 a.a.) soluble protein that is subject to several posttranslational modifications. It is co-translationally myristoylated at its N terminus (13) and autophosphorylated at several positions (14).

Despite its prime importance in physiology and disease, little is known about the molecular details of the CFTR/PKA-C interaction. The CFTR R domain is largely unstructured (15), and may adopt two major conformations that are at equilibrium with each other. In its “wedged-in” form it is intercalated between the two NBDs preventing NBD dimerization and pore opening ((16, 17); Fig. 1A, states 1 and 6), in its “released” form ((18, 19); Fig. 1A, states 2-5) it allows NBD dimerization and pore opening in response to ATP binding. Whereas for the unphosphorylated R domain the equilibrium is heavily biased towards wedged-in (state 1 vs. 2), for the phosphorylated R domain it is shifted towards released (state 5 vs. 6). Thus, phosphorylation causes “irreversible” CFTR stimulation, terminated only through the action of phosphatases. In addition, simple binding of PKA-C to CFTR can also keep the R domain released, causing additional, “rapidly reversible” stimulation for both unphosphorylated and phosphorylated CFTR ((20); Fig. 1A, states 3-4). However, the structural interactions that underlie activation of CFTR by PKA-C are unknown. In the present study we identify key molecular determinants of the CFTR/PKA-C interaction required for reversible and irreversible CFTR stimulation, respectively.

## Results

### Two distinct regions of the PKA-C substrate binding cleft mediate reversible and irreversible CFTR stimulation

Reversible activation of CFTR by PKA-C binding must reflect stabilization of a released R domain conformation through an interaction between CFTR and PKA-C, but which region of the kinase is involved in this process is unknown. One possibility is that PKA-C uses its substrate binding site to directly “grab” a substrate loop of the R domain. Alternatively, interaction between other parts of the channel and the kinase may promote R-domain release. To differentiate between these possibilities, we studied how reversible and irreversible CFTR current activation is affected by two versions of the PKA inhibitory peptide (PKI), a shorter and a longer segment of the heat-stable protein inhibitor of PKA-C (21). PKI peptides are pseudosubstrates that bind into the PKA-C substrate binding cleft (compare Fig. 1B *left* vs. *right*), but in which the target serine is replaced by an alanine (Fig. 1B, *red*). In the 17-mer PKI(6–22)amide the pseudosubstrate site (Fig. 1B, *right*, *light purple*) is preceded by an amphipathic α- helix (*dark purple*); lack of that helix in the shorter PKI(14–22)amide is reflected by its lower affinity for PKA-C (K_I_ ∼36 nM vs. ∼1.7 nM (22)).

To test competitive inhibition by PKI peptides, we purified bovine PKA-C (bPKA-C) from beef heart ((23); Materials and Methods; Fig. S1A, C), and assessed block of phosphorylation by bPKA-C of the fluorescently labeled soluble peptide substrate TAMRA-kemptide (24). Kemptide phosphorylation is reported by a decrease in mobility on thin layer chromatography (TLC) sheets (20) (Fig. 1D-F, *bottom*), allowing reconstruction of phosphorylation time courses by densitometry (Fig. 1D-F, *top*). In control experiments using 5 nM bPKA-C and 20 μM TAMRA-kemptide ∼50% of the substrate (i.e., ∼10 μM) was phosphorylated in <3 minutes (Fig. 1D), yielding a k_cat_ estimate of ∼12 s^-1^ (Fig. 1C, *red bar*). In rough agreement with the reported K_I_ values and the K_M_ of TAMRA-kemptide (∼2 μM (24)), application of 0.2 μM PKI(6–22)amide (Fig. 1E) or 10 μM PKI(14–22)amide (Fig. 1F) slowed phosphorylation time courses by 7-8-fold (Fig. 1C, *striped bars*).

Exposure of inside-out macropatches containing wild-type (WT) CFTR channels to 2 mM ATP + 300 nM bPKA-C (Fig. 1G, *gray* and *red bars*) causes both reversible and irreversible CFTR current activation. The irreversible component which survives bPKA-C removal (Fig. 1G; Fig. 1L, *gray bars*) reports the effect on channel gating of phosphorylation *per se*, and develops over the time course of tens of seconds. Applying bPKA-C first for 1 minute, and then for additional 3 minutes to reach full phosphorylation (Fig. 1G), reveals that the irreversible component reaches ∼70% of its final value already after 1 minute in bPKA-C (Fig. 1L, compare *gray bars* identified by *double arrow*), whereas the rapidly reversible component roughly doubles the open probability of phosphorylated channels (Fig. 1L, *red bars*). Co-application of 5 μM PKI(6–22)amide during the first 1-minute exposure to bPKA-C (Fig. 1H; 1st *purple bar*) abolished current activation, reporting complete inhibition of channel phosphorylation (Fig. 1M, compare *arrowed gray bars*). Moreover, reversible stimulation of phosphorylated channels by re-applied bPKA-C (Fig. 1H; 3rd *red bar*) was also entirely blocked by co-applied PKI(6–22)amide (Fig. 1H; 2nd *purple bar*; Fig. 1M, *striped* vs. *red bar*). When 20 μM PKI(14–22)amide was co-applied during the first 1-minute bPKA-C exposure (Fig. 1I, 1st *light purple bar*) irreversible current activation was also greatly reduced, reaching only ∼8% of its final value (Fig. 1N, compare *arrowed gray bars*). However, in stark contrast to the action of the longer PKI, the shorter peptide failed to suppress rapid reversible stimulation of phosphorylated channels by re-applied bPKA-C (Fig. 1I; 3rd *red bar*; Fig. 1N, *striped* vs. *red bar*). A systematic screening of PKI peptides with the N-terminus incrementally extended from position 14 to 6 further narrowed down the region which blocks reversible CFTR activation to PKI residues 6-9 (Fig. S2).

Pure reversible activation of unphosphorylated channels by bPKA-C can be studied in the presence of 10 μM N^6^-(2-phenylethyl)-ATP (P-ATP; Fig. 1J-K, *dark gray bars*), an ATP analog that opens CFTR channels with high affinity but cannot be used by PKA-C for phosphotransfer (20). The strong reversible stimulation of unphosphorylated channels by bPKA-C in P-ATP was also largely prevented by 5 μM PKI(6–22)amide (Fig. 1J, *purple bar*; Fig. 1O, *striped* vs. *red bar*), whereas it was unaffected by PKI(14–22)amide even at a concentration as high as 100 μM (Fig. 1K, *light purple bar*; Fig. 1P, *striped* vs. *red bar*). Thus, the specific subregion of the PKA-C substrate cleft which accommodates the N-terminal helix (residues 6-13) of PKI(6–22)amide (Fig. 1B, *dark purple*) is responsible for rapid reversible stimulation of both unphosphorylated and phosphorylated CFTR (Fig. 1A, states 3-4).

### PKA-C posttranslational modification pattern profoundly affects efficiency of CFTR stimulation

To facilitate structure/function studies of the CFTR/PKA-C interaction, the bovine PKA-C protein was expressed in *E. coli* and affinity-purified to homogeneity (Fig. S1B, C). Although the amino acid sequence of this recombinant PKA-C (rPKA-C) is identical to that of native bPKA-C, their posttranslational modification patterns are different (25). We therefore first compared their catalytic activities on TAMRA-kemptide (Fig. 2A-B). Phosphorylation time courses for parallel preparations of rPKA-C and bPKA-C were similar, reporting k_cat_ values of ∼7 s^-1^ and ∼9 s^-1^, respectively (Fig. 2C; p=0.085).

**Figure 2.**
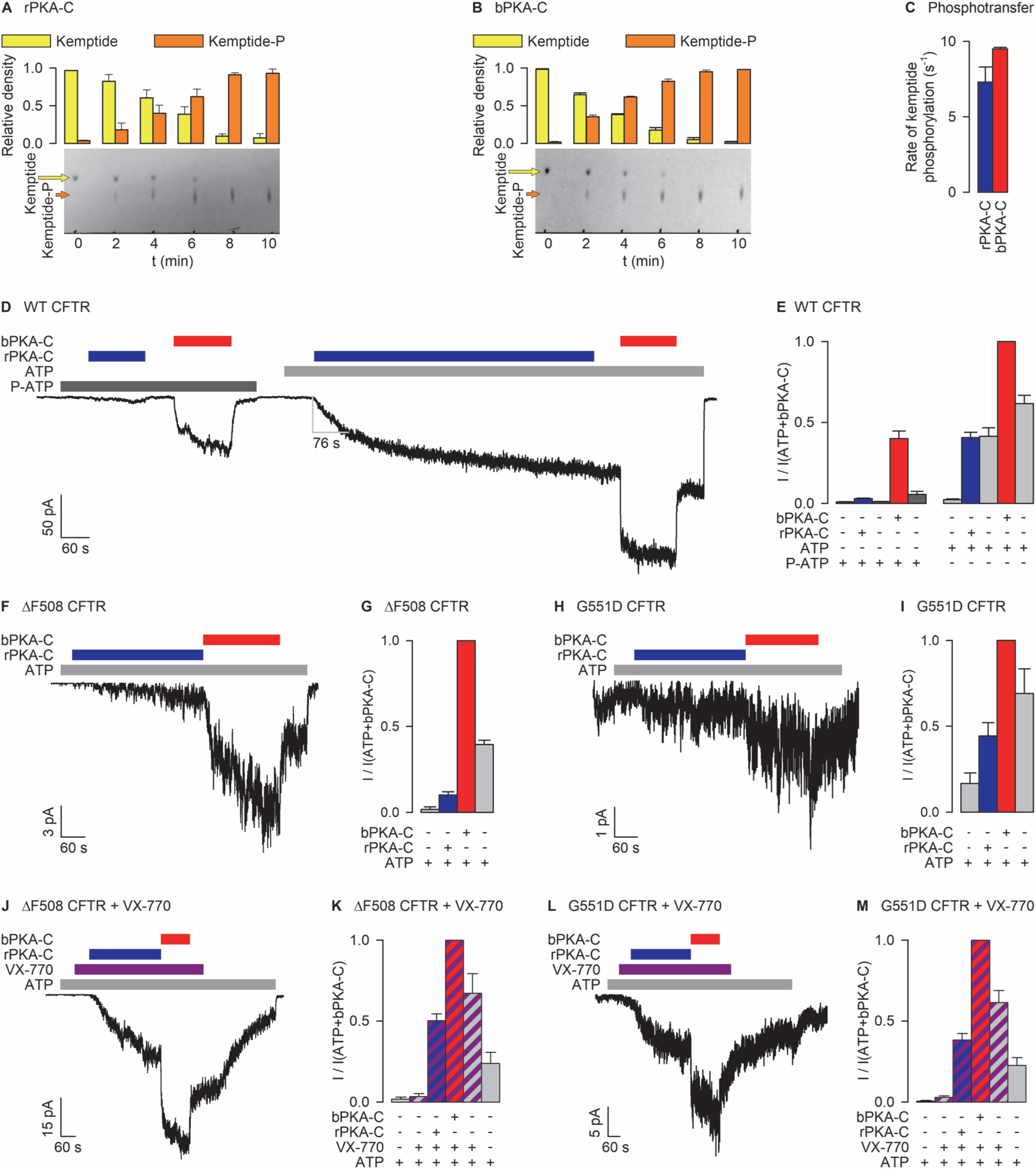
Recombinant PKA-C is fully active on a soluble substrate, but poorly activates CFTR. *A*-*B*, Time courses of kemptide phosphorylation resolved on TLC sheets (*Bottom*) and densitometric analysis (*Top*) for rPKA-C (*A*) or bPKA-C (*B*). Conditions and color coding as in Fig. 1D-F. *C*, Calculated *k*_cat_ (s^-1^) for rPKA-C (*blue*) and bPKA-C (*red*). Data in *A*-*C* represent mean±SEM, n=3. *D*, Macroscopic WT CFTR current activated by repeated exposures to 300 nM rPKA-C (*blue bars*) or bPKA-C (*red bars*), in the presence of either 10 μM P-ATP (*dark gray bar*) or 2 mM ATP (*light gray bar*). *E*, Steady-state currents in the sequential segments of the experiment in *D*, normalized to that in the presence of ATP+bPKA-C (*2nd red column*). Data represent mean±SEM, n=5-6. *F*, *H*, *J*, *L*, Macroscopic currents from ΔF508 (*F*, *J*) or G551D (*H*, *L*) CFTR channels activated by 300 nM rPKA-C (*blue bars*) or bPKA-C (*red bars*) in 2 mM ATP (*gray bars*), in the absence (*F*, *H*) or presence (*J*, *L*) of 10 nM VX-770 (*purple bars*). *G*, *I*, *K*, *M*, Steady-state currents in the sequential segments of the experiment in the preceding panel, normalized to that in the presence of ATP+bPKA-C (*G*, *I*) or ATP+bPKA-C+VX-770 (*K*, *M*). Data represent mean±SEM, n=4-6.

Surprisingly, despite being fully catalytically competent on a soluble substrate, CFTR current activation by rPKA-C was greatly impaired compared to that by bPKA-C (Fig. 2D). In the presence of P-ATP 300 nM rPKA-C failed to reversibly activate unphosphorylated CFTR (Fig. 2D-E, *left*, *blue bars*). In the presence of ATP (Fig. 2D, *right*) CFTR current activation by rPKA-C was greatly slowed compared to that by bPKA-C (cf., Fig. 1G), and the maximal current achieved after ∼5 minutes was less than half of that evoked by subsequent exposure to 300 nM bPKA-C (Fig. 2D-E, *right*, *blue* vs. *red bar*). That smaller maximal current was in part due to a complete lack of reversible stimulation of phosphorylated channels by rPKA-C, as evidenced by a lack of current decline following its removal (Fig. 2E, *right*, *2^nd^* vs. *3^rd^ bar*). In addition, the steady current that survived rPKA-C removal was significantly (p=0.012) smaller than the current following a subsequent exposure to bPKA-C (Fig. 2E, *right*, *3^rd^* vs. *5^th^ bar*), suggesting that CFTR channels are phosphorylated by rPKA-C not only far slower, but also to a somewhat lower final stoichiometry. Similar results were obtained when comparing PKA-C proteins from various commercial sources: whereas native preparations (Sigma-Aldrich P2645 and 539576) functionally resembled our bPKA-C, recombinant products (Promega V5161) recapitulated our findings on rPKA-C. These results suggest that the CFTR/PKA-C interaction is highly sensitive to some posttranslational modification of PKA-C.

Because PKA-C posttranslational modification patterns might dynamically change *in vivo*, differential efficacies of different PKA-C isoforms might bear relevance to CF disease. We therefore compared the efficacies of rPKA-C and bPKA-C for activation of CFTR channels bearing the ΔF508 or G551D mutation. Interestingly, rPKA-C was even less efficient in activating currents for the disease mutants. Compared to bPKA-C, maximal current activated by rPKA-C was only ∼10% and 40%, respectively, for ΔF508 and G551D CFTR (Fig. 2F, H; Fig. 2G, I, *blue* vs. *red bars*).

An increasing number of CF patients worldwide receive treatment with the potentiator VX-770, administered alone or in combination with corrector drugs. As CFTR gating stimulation by VX-770 requires prior phosphorylation of CFTR (3), and rPKA-C might fail to fully phosphorylate mutant CFTR channels (Fig. 2F-I), we compared the efficacies of rPKA-C and bPKA-C for channel activation in the presence of the drug (Fig. 2J, L). As expected, unphosphorylated channels were only marginally stimulated by 10 nM VX-770 (Fig. 2K, M; *2^nd^ bar*), a quasi-saturating concentration of the drug (EC_50_∼1.5 nM, (26)). In addition, even in the presence of the drug, for both mutants steady-state activation by rPKA-C was substantially reduced compared to that by bPKA-C (Fig. 2K, M; compare *3^rd^* and *4^th^ bars*).

### Enzymatic demyristoylation of bovine PKA-C diminishes reversible and irreversible CFTR activation

One documented structural difference between the two PKA-C isoforms is that native bPKA-C is N-myristoylated (13) whereas rPKA-C is not (27). To address whether the PKA-C N-myristoyl group is essential for a fruitful CFTR/PKA-C interaction, we took advantage of the Invasion plasmid antigen J (IpaJ) protein, a cysteine-protease injected by *Shigella flexneri* into the cytosol of infected cells. When expressed *in vitro*, the IpaJ catalytic core domain (residues 51-259) non-specifically demyristoylates a large number of proteins, including PKA-C, by cleaving the peptide bond following the N-myristoylated glycine (28). The IpaJ catalytic domain was expressed in *E. coli* and affinity-purified (Fig. S3A-B). To test its activity, a fluorescently labeled synthetic peptide encompassing the first 11 residues of PKA-C was synthesized in both N-myristoylated (“myr-peptide”) and non-myristoylated (“nonmyr-peptide”) forms. The higher mobility of the myr-peptide on TLC sheets (Fig. 3A, *lanes 1* and *2*) allows to monitor its demyristoylation. Indeed, a 10-minute incubation of 50 μM myr-peptide with 3 μM IpaJ at room temperature completely converted the myr-peptide into a low-mobility product, consistent with cleavage of the N-terminal myr-glycine group (Fig. 3A, *lane 3*).

**Figure 3.**
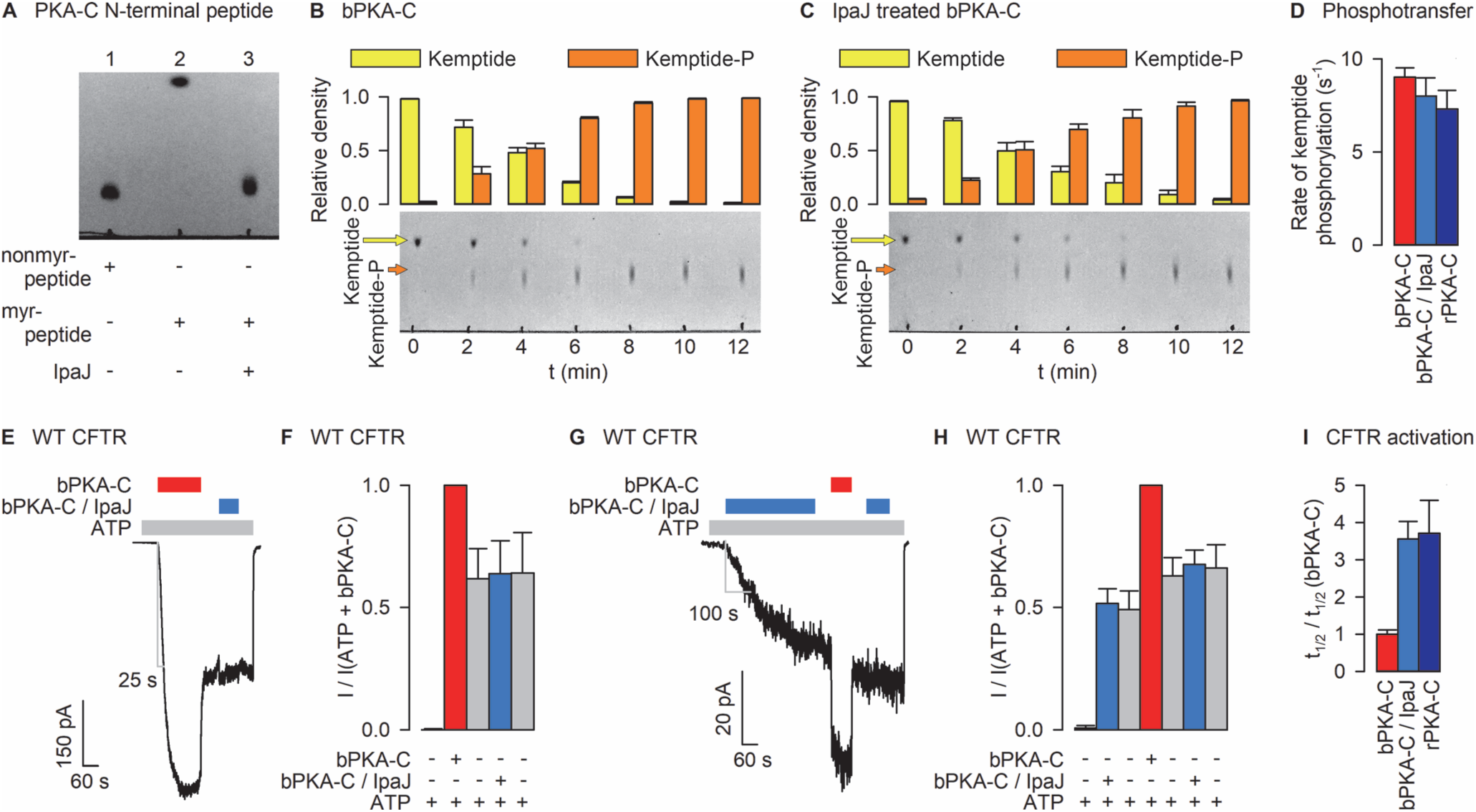
Enzymatic de-myristoylation of bPKA-C selectively impairs efficiency towards CFTR. *A*, TLC sheet showing de-myristoylation of synthetic PKA-C N-terminal peptide by purified IpaJ. Nonmyr-peptide, GNAAAAKKGSE-K-ε-TAMRA (*lane 1*); myr-peptide, N-myristoyl-GNAAAAKKGSE-K-ε-TAMRA (*lane 2*); 50 μM myr-peptide was incubated with 3 μM IpaJ for 10 minutes at room temperature (*lane 3*). *B*-*C*; Time courses of kemptide phosphorylation resolved on TLC sheets (*Bottom*) and densitometric analysis (*Top*) for control bPKA-C (*B*) or IpaJ-treated (see Materials and Methods) bPKA-C (*C*). Conditions and color coding as in Fig. 1D-F. *D*, Calculated *k*_cat_ (s^-1^) for bPKA-C (*red*), IpaJ-treated bPKA-C (*light blue*), and rPKA-C (*dark blue*; replotted from Fig. 2C). Data in *B*-*D* represent mean±SEM, n=3. *E*, WT CFTR current initially activated by 300 nM bPKA-C (*red bar*) and then exposed to 300 nM IpaJ-treated bPKA-C (*light blue bar*). *G*, WT CFTR current initially activated by 300 nM IpaJ-treated bPKA-C (*light blue bar*) and then exposed to 300 nM bPKA-C (*red bar*). *F*, *H*, Steady-state currents in the sequential segments of the experiment in the preceding panel, normalized to that in the presence of ATP+bPKA-C (*red bar*). Data represent mean±SEM, n=7 (*F*) and 5 (*H*). *I*, Time required for half-maximal CFTR current activation (t_1/2_; cf., gray L-bars in Fig. 2D, 3E, 3G) for IpaJ-treated bPKA-C (*light blue bar*) and rPKA-C (*dark blue bar*), normalized to t_1/2_ for activation by bPKA-C obtained in patches from the same batches of oocytes (*red bar*). Data represent mean±SEM, n=5-10.

To test the effect of IpaJ treatment on full-length bPKA-C, the kinase was purified from beef heart, tested for catalytic activity on kemptide (Fig. 3B; Fig. 3D, *red bar*), and then incubated with IpaJ (molar ratio 5:1) at 37°C for 3 hours. IpaJ was then selectively removed and bPKA-C concentration redetermined (Materials and Methods; Fig. S3C). IpaJ treatment did not affect the catalytic activity of bPKA-C towards soluble kemptide (Fig. 3C; Fig. 3D, *light blue bar*). However, WT CFTR channels pre-phosphorylated by a 3-minute exposure to 300 nM untreated bPKA-C (Fig. 3E, *red bar*) failed to respond to subsequent application of 300 nM IpaJ-treated bPKA-C (Fig. 3E, *light blue bar*), revealing complete lack of reversible stimulation of phosphorylated CFTR (Fig. 3F, *red* vs. *light blue bar*). Exposure of unphosphorylated WT CFTR channels to 300 nM IpaJ-treated bPKA-C resulted in slower current activation compared to untreated bPKA-C (cf., Figs. 3E, G), reporting a slowed rate of CFTR phosphorylation, and failed to maximally activate channel currents (Fig. 3G-H, *first blue* vs. *red bar*). To quantitatively compare rates of CFTR current activation by PKA-C variants applied at 300 nM, the time required for half-maximal current activation (t_1/2_; cf., *gray L-bars* in Figs. 2D, 3E, 3G) was normalized to t_1/2_ for activation by bPKA-C measured in the same batches of cells. Just as for rPKA-C (Fig. 3I, *dark blue bar*), t_1/2_ for IpaJ-treated bPKA-C was ∼4-fold longer compared to untreated bPKA-C (Fig. 3I, *red* vs. *light blue bar*). Thus, IpaJ-treated bPKA-C activates CFTR channels as inefficiently as rPKA-C.

### Enzymatic myristoylation of recombinant PKA-C partially restores efficiency of CFTR activation

N-myristoyl transferase (NMT) is not endogenously present in *E. coli*, but recombinant PKA-C can be myristoylated either by co-expression of NMT with PKA-C in the bacteria (29), or by *in vitro* co-incubation of purified rPKA-C and NMT proteins with myristoyl-CoA (30). *In vitro* incubation of 100 μM nonmyr-peptide with 17 μM purified NMT protein + 100 μM myr-CoA indeed resulted in the appearance of the myr-peptide spot (Fig. 4A, lane 3) which could be readily reconverted into the slowly migrating form by subsequent treatment with IpaJ (Fig. 4A, lane 4). However, *in vitro* myristoylation remained incomplete (fractional myristoylation failed to reach 100% even following overnight incubation; Fig. 4A, lane 3), and attempts to *in vitro* myristoylate rPKA-C resulted in protein aggregation.

**Figure 4.**
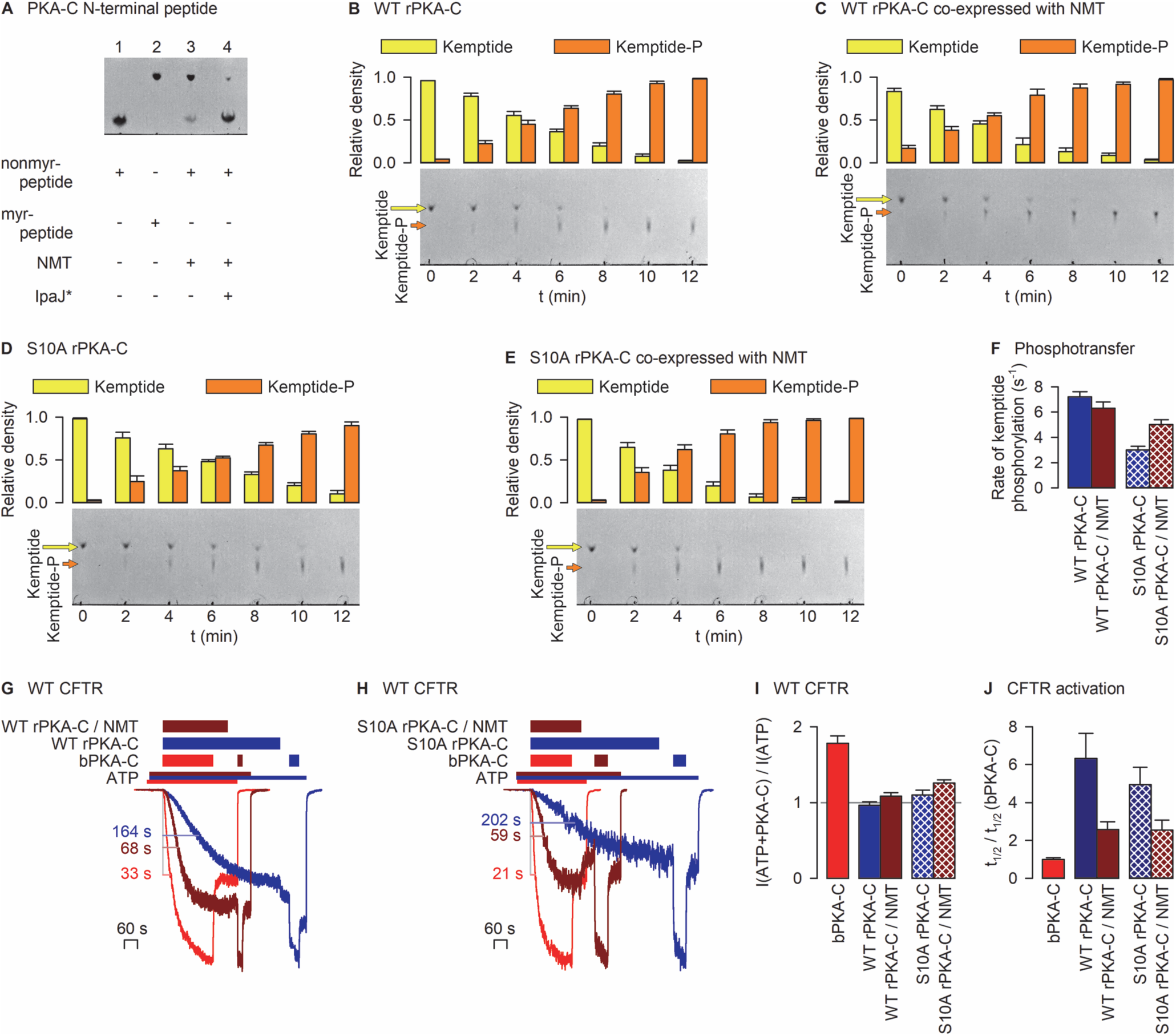
NMT co-expression partially restores rPKA-C efficiency towards CFTR. *A*, TLC sheet showing *in vitro* myristoylation of synthetic PKA-C N-terminal peptide by purified NMT. Controls, nonmyr-peptide (*lane 1*) and myr-peptide (*lane 2*); 100 μM nonmyr-peptide was incubated with 17 μM NMT + 100 μM myr-CoA overnight at room temperature (*lane 3*). *An aliquot of the reaction product was subsequently treated with 3 μM IpaJ for 10 minutes (*lane 4*). *B*-*E*; Time courses of kemptide phosphorylation resolved on TLC sheets (*Bottom*) and densitometric analysis (*Top*) for control rPKA-C (*B*), rPKA-C co-expressed with NMT (*C*), control S10A rPKA-C (*D*), or S10A rPKA-C co-expressed with NMT (*E*). 5 nM rPKA-C (*B*), 8 nM rPKA-C/NMT (*C*), 10 nM S10A rPKA-C (*D*), or 10 nM S10A rPKA-C/NMT (*E*) were incubated with 20 μM TAMRA-kemptide + 200 μM MgATP for the indicated amounts of time at room temperature. Color coding as in Fig. 1D-F. *F*, Calculated *k*_cat_ (s^-1^) for control rPKA-C (*dark blue*), rPKA-C/NMT (*brown*), control S10A rPKA-C (*dark blue checkered*), and S10A rPKA-C/NMT (*brown checkered*). Data in *B*-*F* represent mean±SEM, n=3. *G*-*H*, Overlays of three WT CFTR current traces, activated by 300 nM of (*G*) bPKA-C (*red trace*), rPKA-C (*dark blue trace*), or rPKA-C/NMT (*brown trace*) and (*H*) bPKA-C (*red trace*), S10A rPKA-C (*dark blue trace*), or S10A rPKA-C/NMT (*brown trace*), respectively. For the *dark blue* and *brown traces* 300 nM bPKA-C was subsequently applied. Currents were synchronized to the time point of first PKA-C application, and normalized to the steady-state amplitude in 300 nM bPKA-C. Color coding of bars that identify exposure to compounds follows that of the current traces. *I*, Reversible stimulation by bPKA-C (*red*), rPKA-C (*dark blue*), rPKA-C/NMT (*brown*), S10A rPKA-C (*dark blue checkered*), and S10A rPKA-C/NMT (*brown checkered*), expressed as the ratio of mean steady current in ATP+PKA-C to that following PKA-C removal. *J*, Time required for half-maximal CFTR current activation (t_1/2_; cf., colored L-bars in panels *G*-*H*) by PKA-C variants, normalized to t_1/2_ for activation by bPKA-C obtained in patches from the respective batches of oocytes. Color coding as in panel *I*. Data in *I*-*J* represent mean±SEM, n=6-12.

To overcome this limitation, PKA-C was co-expressed with NMT in *E. coli*, and rPKA-C was affinity purified (Fig. S1B-C). Phosphotransfer activities of the resulting enzyme (“rPKA-C/NMT”) and of a control preparation of rPKA-C towards kemptide (Fig. 4B-C) were similar (Fig. 4F, *left*). However, a comparison of CFTR current activation kinetics (Fig. 4G) by 300 nM bPKA-C (*red trace*), rPKA-C (*blue trace*) or rPKA-C/NMT (*brown trace*), studied in the same batch of CFTR-expressing cells, revealed a clearly intermediate phenotype for rPKA-C/NMT. Whereas for rPKA-C t_1/2_ for CFTR activation was ∼6-fold longer compared to bPKA-C (Fig. 4J, *left*, *blue* vs. *red bar*), for rPKA-C/NMT it was only ∼2-fold longer (Fig. 4J, *left*, *brown* vs. *red bar*). Consistent with earlier reports (31), these results suggest that NMT co-expression yields partially myristoylated rPKA-C. Based on the effect of NMT co-expression on t_1/2_, the estimated myristoylation efficiency is ∼30%, too low to restore reversible CFTR activation as reflected by a lack of significant current reduction following rPKA- C/NMT removal (Fig. 4I, *brown bar*) (cf., (20)).

### Phosphorylation at serine 10 of PKA-C does not affect its efficiency to activate CFTR

A second distinguishing structural feature of rPKA-C is its reported autophosphorylation at serine 10 (25). Indeed, LC-MS analysis of our PKA-C preparations confirmed that bPKA-C is not phosphorylated at this position whereas rPKA-C is phosphorylated to a stoichiometry of ∼60% (5 out of 8 recovered peptide fragments). To address the impact of that difference on CFTR channel activation, PKA-C bearing the S10A mutation was expressed in *E. coli* with or without NMT. As reported previously (32), the majority of the mutant PKA-C was targeted to inclusion bodies, reporting a negative effect of the S10A mutation on folding efficiency. Nevertheless, for both S10A rPKA-C and S10A rPKA-C/NMT sufficient amounts could be purified from the soluble fractions (Fig. S1B, C) to allow functional analysis. Both mutant proteins were active when assayed on kemptide (Fig. 4D-E), but showed a modest reduction in phosphotransfer rates (Fig. 4F, cf., *checkered* vs. *non-checkered bars*) which was significant for S10A rPKA-C (∼2-fold, p=0.0005). However, CFTR channel activation by S10A rPKA-C (Fig. 4H, *blue trace*) resembled that by WT rPKA-C: t_1/2_ for current activation was ∼5- fold prolonged compared to bPKA-C (Fig. 4J, *blue checkered bar*), and reversible stimulation was undetectable (Fig. 4I, *blue checkered bar*). Importantly, just as for WT rPKA-C, NMT co-expression increased the efficiency of CFTR current activation by S10A rPKA-C (Fig. 4H, *brown trace*): t_1/2_ was shortened (Fig. 4J, *brown checkered bar*), and even a small (∼1.25-fold) but significant (p=0.003) reversible stimulation was restored (Fig. 4I, *brown checkered bar*).

### Uncharacterized IpaJ homologs are present in the genomes of Gram-negative CF pathogens

*Pseudomonas*, *Burkholderia*, and *Stenotrophomonas* genera are among the most common CF pathogens, and all three contain secretion systems that deliver effector proteins into the host cell cytosol (33). However, to date only a small subset of these secreted virulence factors have been identified. The *Shigella* effector protein IpaJ is a member of the cysteine peptidase C39-like family which includes many uncharacterized members in more than 200 bacterial species (28, 34). Using a BLAST search we identified several IpaJ homologs also in the three most CF-relevant bacterial genera (Fig. 5A).

**Figure 5.**
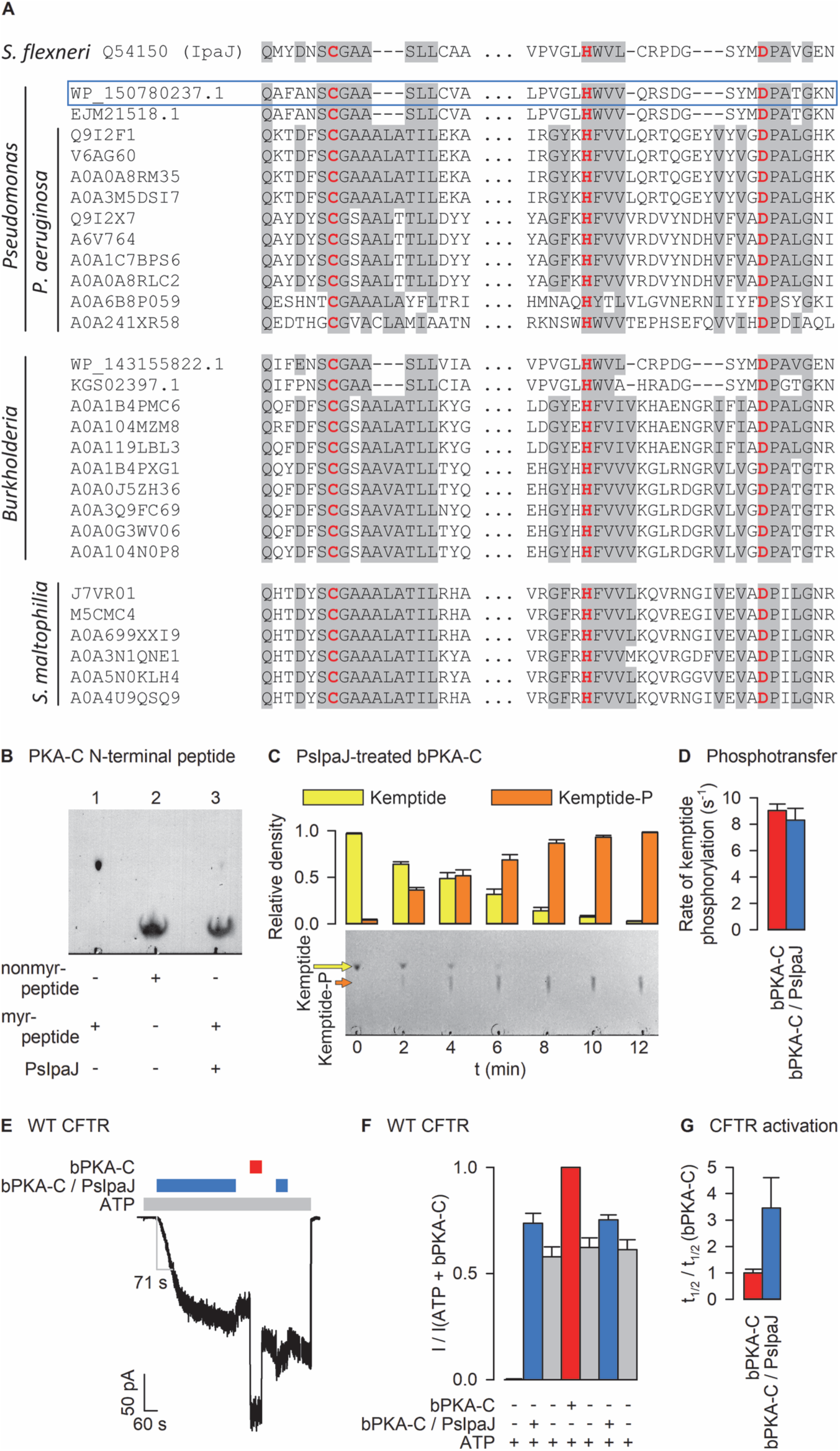
Pseudomonas IpaJ homolog de-myristoylates bPKA-C *in vitro*. *A*, Sequence alignment of catalytically important segments of *Shigella flexneri* IpaJ with homologous sequences identified in the genomes of *Pseudomonas* (top block), *Burkholderia* (center block) and *Stenotrophomonas* (bottom block) genera. *Gray*, conserved residues; *bold red*, cysteine protease catalytic triad; *blue box*, *Pseudomonas* sequence most closely related to IpaJ (PsIpaJ). *B*, TLC sheet showing de-myristoylation of synthetic PKA-C N-terminal peptide by purified PsIpaJ. Myr-peptide, N-myristoyl- GNAAAAKKGSE-K-ε-TAMRA (*lane 1*); nonmyr-peptide, GNAAAAKKGSE-K-ε-TAMRA (*lane 2*); 30 μM myr-peptide was incubated with 0.5 μM PsIpaJ for 30 minutes at room temperature (*lane 3*). *C*; Time course of kemptide phosphorylation resolved on TLC sheet (*Bottom*) and densitometric analysis (*Top*) for PsIpaJ-treated (see Materials and Methods) bPKA-C. Conditions and color coding as in Fig. 1D-F. *D*, Calculated *k*_cat_ (s^-1^) for bPKA-C (*red*; replotted from Fig. 3D) and PsIpaJ-treated bPKA-C (*light blue*) from the densitometric analysis in *C*. Data represent mean±SEM, n=3. *E*, WT CFTR current initially activated by 300 nM PsIpaJ-treated bPKA-C (*light blue bar*) and then exposed to 300 nM untreated bPKA-C (*red bar*). *F*, Steady-state currents in the sequential segments of the experiment in *E*, normalized to that in the presence of ATP+bPKA-C (*red bar*). *G*, Time required for half-maximal CFTR current activation (t_1/2_; cf., gray L-bar in panel *E*) for PsIpaJ-treated bPKA-C (*light blue bar*), normalized to t_1/2_ for activation by bPKA-C obtained in patches from the same batches of oocytes (*red bar*). Data in panels *F*-*G* represent mean±SEM, n=8-9.

To test whether enzymes capable of removing N-myristoyl glycines from target proteins may be present also in these bacteria, we expressed and purified (Fig. S3A-B) the catalytic domain of the *Pseudomonas* sequence with highest homology to IpaJ (WP_150780237.1; Fig. 5A, *blue frame*). When tested on the PKA-C N-terminal myr-peptide, this “PsIpaJ” protein also readily removed the N-myristoyl glycine (Fig. 5B). Moreover, treatment of bPKA-C with PsIpaJ (Fig. S3D) resulted in functional effects similar to those of IpaJ treatment. Compared to untreated bPKA-C, CFTR current activation by PsIpaJ-treated bPKA-C was ∼3-fold slower (Fig. 5E; Fig. 5G, *light blue bar*), and reversible CFTR activation was largely diminished, as reported by a virtual lack of current decline upon its removal (Fig. 5E, Fig. 5F *blue* vs. subsequent *gray bars*). On the other hand, the k_cat_ for kemptide phosphorylation was unaffected by PsIpaJ treatment (Fig. 5C-D).

## Discussion

Despite its relevance for physiological regulation, CF disease, cholera, and ADPKD, the molecular mechanisms by which PKA-C activates CFTR have only started to become clear (Fig. 1A). PKA-C binding reversibly stimulates both unphosphorylated and phosphorylated CFTR channels by preventing the R domain from returning into its wedged-in position (20). But whether that is achieved through direct binding of R-domain substrate loops by the PKA-C substrate binding site or reflects an allosteric effect triggered by an interaction between other parts of the two proteins is unknown. Here we show that reversible CFTR activation requires a specific small region near to, but distinct from the PKA-C substrate binding site: whereas, as expected, irreversible CFTR stimulation (i.e., phosphorylation) is prevented by both short and long PKI pseudosubstrate peptides (Fig. 1H-I, Fig. 1M-N, *arrowed gray bars*), reversible activation of both unphosphorylated and phosphorylated CFTR channels is specifically blocked by the short amphipathic helix (Fig. 1B, right, *dark purple*) formed by residues 6-13 of PKI(6–22)amide (Fig. 1H-I, Fig. 1M-N, *striped bars*). Comparison of K_I_ values of the short and long peptide suggests a low (millimolar) affinity for binding of the N-terminal helix *per se* to PKA-C. Nevertheless, at a concentration of 1 mM, near its solubility limit, PKI(6–13)amide significantly (p=7.3·10^-5^) suppressed reversible activation of phosphorylated CFTR channels by 300 nM bPKA-C (Fig. S4).

One interpretation is that the small hydrophobic patch on the PKA-C surface which accommodates PKI(6–13)amide serves as a “docking site” (Fig. 6, *top left*, “d”): binding of an R-domain segment adjacent to a target serine loop would help to present the latter for catalysis (Fig. 6, *top left*, *asterisk* marks binding site of target serine). That mechanism would explain why reversible stimulation is preserved even in phosphorylated channels (Fig. 1L, *right red bar*), even though phosphorylation dramatically decreases the binding affinity of target serine loops for the kinase (35). By blocking the entire substrate binding cleft, including both the active site and the “docking site”, the long peptide PKI(6–22)amide would prevent R-domain docking and thereby even reversible stimulation (Fig. 6, *bottom right*). In contrast, the shorter PKI(14–22)amide leaves the docking site unoccupied allowing R-domain docking and reversible stimulation (Fig. 6, *bottom left*). Interestingly, reversible stimulation of phosphorylated channels was even slightly (but significantly, p=0.02) enhanced by the presence of PKI(14–22)amide (Fig. 1I; Fig. 1N, *red* vs. *subsequent striped bar*; cf. Fig. S2). Because binding of long PKI peptides induces closure of the substrate cleft (36–38) we speculate that PKI(14–22) binding may also induce some degree of cleft closure which enhances the binding affinity of the “docking site” for R-domain loops. As an alternative to the “docking site” model, interaction of that hydrophobic patch on PKA-C (Fig. 6, “d”) with some other part of CFTR maybe required for promoting R domain release from its wedged-in position.

**Figure 6.**
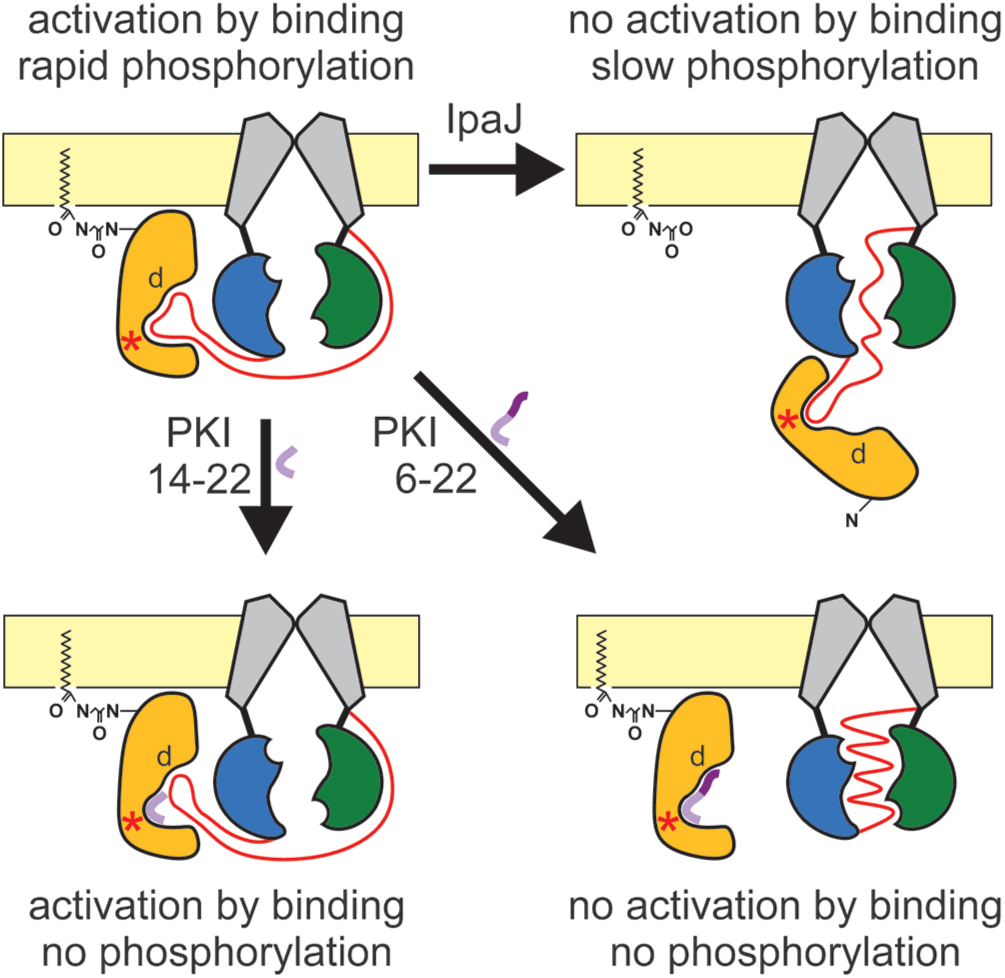
Structural requirements for efficient CFTR/PKA-C interaction. Cartoon color coding as in Fig. 1A. (*Top left*) Membrane-tethering of PKA-C properly orients its catalytic cleft for binding substrate segments of the CFTR R domain, causing both reversible activation and rapid phosphorylation of CFTR. *Red asterisk*, binding site for target serine, “d”, “docking site”. (*Top right*) Cysteine protease-mediated cleavage of the N-myristoyl-glycine from PKA-C untethers the enzyme from the membrane, preventing reversible activation and slowing phosphorylation of CFTR. (*Bottom left*) Pseudosubstrate peptide PKI(14–22)amide (light *violet ribbon*) blocks active site but leaves “docking site” accessible for R-domain binding, preventing phosphorylation but not reversible activation. (*Bottom right*) Pseudosubstrate peptide PKI(6–22)amide (*violet ribbon*; *light segment*, substrate loop; *dark segment*, amphipathic helix) blocks PKA-C/R-domain interaction, preventing both reversible activation and phosphorylation.

We also show here that phosphotransfer by PKA-C is not the rate limiting step for irreversible CFTR activation. If that were the case, with a k_cat_ of ∼10 s^-1^ (Figs. 1C, 2C, 3D, *red*), all ∼11 target serines in the R domain should be phosphorylated within ∼1 s, whereas CFTR current activation by bPKA-C requires ∼1 minute (Figs. 1G, 3E). Correspondingly, the modest reduction in phosphotransfer rate of S10A rPKA-C (Fig. 4F, *blue checkered* vs. *uncheckered bar*) does not slow CFTR current activation relative to that by WT rPKA-C (Fig. 4J, *blue checkered* vs. *uncheckered bar*). Thus, other kinetic steps, likely related to spontaneous release of the unphosphorylated R domain from its wedged-in position (Fig. 1A, step 1→2), and/or targeting of PKA-C to its unstructured substrate, rate limit CFTR activation.

Along those lines, we discover here that the presence of an N-terminal myristoyl group on PKA-C fundamentally determines its efficiency towards CFTR. Whereas both myristoylated (bPKA-C) and non-myristoylated (rPKA-C) PKA-C are equally effective on a soluble substrate (Figs. 2A-C, 3B-D), non-myristoylated PKA-C only poorly activates CFTR. In particular, 300 nM non-myristoylated PKA-C supports no reversible activation (Fig. 2D-E) and ∼4-6-fold slower irreversible activation compared to 300 nM myristoylated PKA-C (Fig. 3I, Fig. 4J, *blue*). Moreover, even at steady state, full phosphorylation is not reached within the entire time course of our experiments (Fig. 2D-E), suggesting that some R-domain serines might not become phosphorylated. Intriguingly, that inefficiency of non-myristoylated PKA-C is even more pronounced towards disease mutants ΔF508 and G551D for which maximal current in rPKA-C is only ∼10% and ∼40%, respectively, of that in bPKA-C (Fig. 2F-I).

In support of this crucial role of the PKA-C N-myristoyl group, its enzymatic removal from the native enzyme diminishes (Fig. 3E-I), whereas its incorporation into the recombinant enzyme enhances (Fig. 4G-J), efficiency towards CFTR. In contrast, neither of these manipulations affect phosphotransfer activity towards a soluble substrate (Figs. 3B-D, 4B-F). The alternative explanation that the presence of phosphoserine 10 in rPKA-C impairs its efficiency towards CFTR is ruled out by the lack of effect of the S10A mutation on CFTR activation time courses (Fig. 4G-H, *blue traces*; Fig. 4J, *blue checkered* vs. *non-checkered bars*).

Co-expression of NMT with PKA-C in *E. coli* did not result in fully myristoylated rPKA-C (Fig. 4G-J). One possible reason for that inefficiency is the reported autophosphorylation of S10 in rPKA-C (25), which impedes N-myristoylation (31). However, although NMT co-expression restored a significant fraction of reversible stimulation for S10A (but not WT) rPKA-C (Fig. 4I), its limited amplitude suggests that myristoylation stoichiometry remained incomplete even for the mutant. Thus, other factors, perhaps the availability of myristoyl-CoA in the bacteria, must be limiting.

The presence of an N-myristoyl group on PKA-C was discovered decades ago (13), but its exact biophysical role is still not entirely clear. Although N-myristoylated, PKA-C is a soluble cytosolic protein (39), and the N-myristoyl group is not required for its association with the regulatory subunit, cAMP-dependent activation, phosphotransfer activity towards most substrates, or access to the nucleus (40, 41). The Type II holoenzyme is targeted to membranes, but this is thought to be primarily achieved through binding of the Type II regulatory subunit (RII) to membrane-associated A Kinase Anchoring Proteins (AKAPs (42)). More recent work revealed that a rapid equilibrium of the PKA-C N-myristoyl group between solvent exposed (myr-out) and protein-occluded (myr-in) conformations (43) is shifted towards myr-out by interaction with membranes (30) or with RII-type regulatory subunits (44). Moreover, the PKA-C N-myristoyl group allows AKAP-independent membrane anchoring of Type II holoenzymes (45) and also of free PKA-C subunits (46). Indeed, in intact cells, stimulation of the cAMP pathway resulted in membrane accumulation of predominantly free PKA-C subunits, allowing preferential phosphorylation of membrane substrates (46). To our knowledge the present study on CFTR is the first to identify a membrane substrate which can efficiently interact exclusively with myristoylated PKA-C. That specificity is likely based on both proximity and orientation effects. Membrane accumulation increases the local concentration of myristoylated PKA-C near the CFTR channel, and physical attachment to the membrane restrains PKA-C in an optimal orientation, required for engaging the CFTR R domain (Fig. 6, *top left*). Severing the membrane tether from PKA-C eliminates reversible stimulation and impairs R-domain phosphorylation (Fig. 6, *top right*).

Beyond its biophysical implications these findings might bear relevance for human disease. Until recently the presence of an N-myristoyl group was considered a permanent feature of a subset of proteins, but this has changed with the discovery of the *Shigella* virulence factor IpaJ, the first enzyme shown to catalyze the removal of N-myristoyl groups from proteins. A major challenge in CF patient care is the treatment of recurrent lung infections, accompanied by exacerbation of pulmonary function. Such infections are most often caused by *Pseudomonas*, *Burkholderia*, and *Stenotrophomonas* species that eventually colonize the CF lung (33). Depending on the stage of the infection, these strains employ Type III, IV, and/or VI secretion systems (33, 47), or outer membrane vesicles (48, 49) to directly deliver virulence factors into the cytosol of infected cells, with or without close contact to them. However, very few of these effector proteins have been studied to date. Here we identify in their genomes a large number of uncharacterized cysteine-protease sequences homologous to IpaJ (Fig. 5A), and show for one *Pseudomonas* protein that it readily demyristoylates PKA-C *in vitro* (Fig. 5B, E-G). Because non-myristoylated PKA-C activates mutant CFTR channels even less efficiently (Fig. 2F-I), also in the presence of a potentiator (Fig. 2J-M), PKA-C demyristoylation in lung epithelial cells might further impair residual CFTR function in infected CF patients. Of note, although IpaJ also demyristoylates a broad range of target proteins *in vitro* (28), in host cell infection models its substrate specificity was narrow suggesting an unknown regulatory mechanism (34). Thus, among the large number of IpaJ homologs in CF pathogens it will be interesting to search for enzymes that target PKA-C also *in vivo*. Identification of such effector proteins might define novel drug targets for the treatment of CF lung disease.

## Acknowledgments

We thank Alexandra Reers and Robin Prentice (Seattle Structural Genomics Center for Infectious Disease) for providing the coding sequence for *Cryptosporidium parvum* NMT and purified NMT protein, and Dr. Angus Nairn (Yale University) for expert advice on bPKA-C preparation. Supported by EU Horizon 2020 Research and Innovation Program grant 739593 and Cystic Fibrosis Foundation Research Grant CSANAD21G0 to L.C., as well as by National Research, Development and Innovation Fund grants KKP 144199 to L.C. and PD 131643 to C.M.

## Author contributions

All authors contributed to designing the project, performing experiments, analyzing and interpreting data, and writing the manuscript.

## Declaration of interests

The authors declare no competing interests.

## Data Availability

Data generated and analyzed over the course of the current study are included within the paper and SI Appendix.

## Materials and Methods

### Xenopus laevis oocyte isolation and injection

Oocytes were isolated from anaesthetized adult female *Xenopus laevis* following Institutional Animal Care Committee guidelines, injected with 10 ng cRNA in a fixed 50-nl volume, and stored at 18°C as described (50). Current recordings were obtained 1-3 days after injection.

### Molecular biology

The coding sequences of full-length bovine protein kinase A catalytic subunit alpha (Genbank accession number: NP_777009.1) and of the catalytic domains of IpaJ (Genbank accession number: Q54150, residues 51-259) and PsIpaJ (Genbank accession number: WP_150780237, residues 29-234), with C-terminal Twin-Strep-tags, were sequence-optimized for expression in *E. coli*, synthesized (General Biosystems) and incorporated into the pJ411 plasmid (DNA2.0) bearing kanamycin resistance.

N-myristoyl transferase (NMT) from *Cryptosporidium parvum* Iowa II (Genbank accession number: Q5CV46) with C-terminal His_6_-tag, subcloned into the pET14b plasmid bearing ampicillin resistance, was a kind gift from Alexandra Reers and Robin Prentice, Seattle Structural Genomics Center for Infectious Disease.

The cDNA for WT and mutant CFTR in pGEMHE was linearized (Nhe I, New England Biolabs), transcribed *in vitro* (mMESSAGE mMACHINE T7 Transcription Kit, ThermoFisher Scientific), and purified cRNA stored at −80 °C.

### Purification of native bovine PKA-C from beef heart

The catalytic subunit of protein kinase A was purified from beef heart as described (23). Approximately 2.7 kg cleaned myocardium was ground and homogenized in 6 liters of Buffer A (10 mM potassium-phosphate (pH=6.9), 1 mM EDTA) supplemented with 15 mM β-mercaptoethanol and 0.1 mM phenylmethylsulfonyl fluoride (“Buffer A+”), and centrifuged at 14,300 x g for 35 min. The supernatant was filtered through glass wool and mixed with 2 liters of DEAE-sepharose (Merck) pre-equilibrated with Buffer A+. The slurry was pH-adjusted to 6.9, and stirred overnight. The resin was dried in a Büchner funnel, washed 4x with 2 liter Buffer A+ and 1x with 2 liter Buffer A+ supplemented with 50 mM NaCl. The dried resin was mixed with 2 liters of Buffer A+ supplemented with 500 mM NaCl (Buffer B+), stirred for 2 h, filtered through a Büchner funnel, and washed with additional 2 liters of Buffer B+. The flowthrough (1st eluate, ∼4.5 liters) was salted out by gradual addition of 390 g/l (NH_4_)_2_SO_4_ under constant stirring followed by centrifugation at 14,300 x g for 35 min. The pellet was resuspended in 450 ml Buffer A+ and dialysed against 8 liters of Buffer A+ for ∼15 h, with Buffer A+ replaced 2x. The dialysed sample (∼600 ml) was centrifuged at 16,000 x g for 30 min, and the pH of the supernatant adjusted to 6.1 with 1M acetic acid. To absorb impurities with pI>6.1, the sample was mixed with 200 ml CM Sephadex C-50 resin preequilibrated with Buffer A+ (pH=6.1), stirred for 5 minutes and filtered through a Büchner funnel. The resin was discarded and the flowthrough was further treated by repeating the absorption protocol 4x at pH=6.1 and then 5x at pH=6.9. The sample was centrifuged at 17,700 x g for 30 min, the supernatant was supplemented with 200 μM cAMP and loaded overnight (ÄKTA Purifier 10) onto a 20-ml CM Sephadex C-50 column preequilibrated with Buffer A+ (pH=6.9). The column was washed with 2x30 ml Buffer A supplemented with 15 mM β-mercaptoethanol, and eluted with a linear salt gradient (50 ml Buffer A vs. 50 ml of 300 mM potassium phosphate (pH=6.9), 1 mM EDTA, 15 mM β-mercaptoethanol). The protein kinase A catalytic subunit eluted at ∼150 mM potassium phosphate (Fig. S1A). Fractions of high purity (>90% by SDS-PAGE; Fig. S1C, *left*) were pooled, and the final concentration determined from the optical density at 280 nm (A_280_; NanoPhotometer P300, Implen GmbH). The entire protocol was carried out at 4-6°C.

### Expression and purification of recombinant bovine PKA-C

pJ411-PKA-C was transformed into *E. coli* BL21(DE3), colonies were grown at 37°C in Luria-Bertani (LB) medium supplemented with 50 μg/ml kanamycin until OD_600_ ∼0.5, and protein expression was induced with 0.1 mM isopropyl-β-D-thiogalactoside (IPTG) overnight at 25°C. Cells were lyzed by sonication in Buffer A (100 mM Tris-HCl (pH=7.5), 150 mM NaCl, 0.1 mM EDTA, 10 mM MgCl_2_, 2 mM DTT) supplemented with Halt Protease Inhibitor Cocktail (Thermo Scientific). Following centrifugation (27,000 x g for 30 min) the cleared supernatant was loaded onto a 5-ml STREP-Tactin Superflow column (IBA Lifesciences) after adding 1.5 mg avidin per liter of starting bacterial culture. Resin-bound PKA-C was eluted with Buffer A supplemented with 10 mM desthiobiotin (IBA Lifesciences). Protein-rich fractions were concentrated (Vivaspin 6, 10,000 MWCO), loaded onto a gel filtration column (Superdex 200 10/300 GL, GE Healthcare) and separated in 20 mM Tris-HCl (pH=7.5), 100 mM NaCl, 10 mM MgCl_2_, 2 mM DTT as the mobile phase (Fig. S1B, *blue chromatograms*). Protein-rich fractions (50-100 µM PKA-C) were collected, pooled, quality-checked by SDS-PAGE (Fig. S1C, *center*), and the final concentration calculated from A_280_.

### Co-expression of Cryptosporidium parvum NMT with recombinant bovine PKA-C in E. coli Rosetta 2(DE3) cells

pJ411-PKA-C and pET14b-NMT were co-transformed into *E. coli* Rosetta 2(DE3) cells Merck (Millipore), colonies were grown at 37°C in LB medium + 50 μg/ml kanamycin, 100 μg/ml ampicillin, and 34 μg/ml chloramphenicol, until OD_600_ ∼0.5, and protein expression was induced with 0.1 mM IPTG + 260 μM Na-myristate overnight at 25°C. Cells were lyzed and PKA-C purified as described above (Fig. S1B, *brown chromatograms*; Fig. S1C, *right*).

### Expression and purification of IpaJ and PsIpaJ

pJ411-IpaJ or pJ411-PsIpaJ was transformed into *E. coli* BL21(DE3), colonies were grown at 37°C in 800 ml of LB medium + 50 μg/ml kanamycin until OD_600_ ∼0.6, and protein expression was induced with 0.5 mM IPTG overnight at 20°C. Cell pellets were resuspended in Buffer A (100 mM Tris-HCl (pH=7.5-8.0), 150 mM NaCl) supplemented with 5% glycerol, 1.5 mM MgCl_2_, 2 mM TCEP and protease inhibitors. Cells were lysed with sonication, and following pellet clearing 1.5 mg/l avidin was added to the soluble fraction before loading on a 5 ml STREP-Tactin Superflow column (IBA Lifesciences) pre-equilibrated with Buffer A + 1 mM TCEP. Following wash with Buffer A + 1 mM TCEP, the protein was eluted with 10 mM desthiobiotin and concentrated (Vivaspin 6, 10,000 MWCO). IpaJ was clean without further purification as verified by SEC (Fig. S3A, *blue chromatogram*), whereas PsIpaJ was further purified by size exclusion chromatography on a Superdex 200 column in a buffer containing 50 mM Tris-HCl (pH=8.0), 150 mM NaCl, 1mM TCEP (Fig. S3A, *red chromatogram*). The quality of both proteins was verified by SDS-PAGE (Fig. S3B), and the final concentrations were calculated from A_280_.

### IpaJ- and PsIpaJ-treatment of native bovine PKA-C

Native bovine PKA-C (in phosphate buffer) was mixed at 20:1 molar ratio with IpaJ (in 100 mM Tris-HCl (pH=7.5), 150 mM NaCl, 1mM TCEP) or at 5:1 molar ratio with PsIpaJ (in 50 mM Tris-HCl (pH=8.0), 150 mM NaCl, 1mM TCEP), and incubated at 37°C for 3 hrs or at 25°C overnight, respectively. The protease was removed by adding STREP-Tactin Sepharose (IBA Lifesciences) resin pre-equilibrated in 50 mM Tris-HCl (pH=8.0), 150 mM NaCl, 1mM TCEP buffer. After a 1-hour rotational mixing of the reaction at 25°C, resin beads were removed by spinning the mixture through Spin-X tubes (Merck) (Fig. S3C-D).

### Kemptide phosphorylation assay

Phosphorylation of the soluble substrate kemptide was done as described (20). PKA-C protein (5-10 nM), TAMRA-kemptide (20 μM; Addexbio Technologies), and ATP (200 μM) were co-incubated at room temperature for 0-12 minutes (or for 0-60 minutes in the presence of PKI peptides) in a reaction buffer containing 50 mM HEPES (pH=7.5), 10 mM Mg-acetate, and freshly added 0.2 mg/mL bovine serum albumin and 5 mM DTT. A 2-μl aliquot of each sample was spotted on a TLC sheet (SIL/UV254, Macherey-Nagel), and the sheet developed in a mixture of n-butanol, pyridine, acetic acid, and water (15:10:3:12 v/v). Kemptide phosphorylation was reported by the appearance of a lower-mobility spot, and the relative amounts of dephospho- vs. phospho-kemptide in each sample were quantitated by densitometry (ImageJ). To estimate k_cat_, the time point of “cross-over” between kemptide and phosphokemptide concentrations (i.e., the time required for phosphorylation of 10 μM kemptide) was calculated by linear time-interpolation between the two bracketing time points. All phosphorylation assays were performed in triplicates. Similar activities were obtained for five different preparations of bPKA-C (k_cat_ ∼8-12 s^-1^) and for three preparations of rPKA-C (k_cat_ ∼7-8 s^-1^) (cf., Figs. 1-5).

### PKA-C N-terminal peptide myristoylation / de-myristoylation assay

*In vitro* myristoylation of PKA-C N-terminal peptide (nonmyr-peptide) was done using an approach described previously (30), with modifications. In brief, final concentrations of 100 µM nonmyr-peptide, 100 µM myristoyl-CoA and 17 µM NMT were added to reaction buffer (100 mM Tris-HCl (pH 7.5), 150 mM NaCl) in a 40-µL reaction mix and left overnight at room temperature. Control reactions did not contain NMT, but were otherwise treated identically.

*In vitro* demyristoylation of myristoylated PKA-C N-terminal peptide (myr-peptide) was done using an approach described previously (28), with modifications. In brief, final concentrations of 50 µM myr-peptide and 3 µM IpaJ was added to reaction buffer (10 mM potassium phosphate (pH 6.9), 1 mM EDTA) in a 40-µL reaction mix and incubated for 10 minutes at room temperature.

Control myr- and nonmyr-peptide, IpaJ-treated myr-peptide, and NMT-treated nonmyr-peptide were spotted (2 µL/spot) on TLC sheets (SIL/UV254, Macherey-Nagel) and separated for about 30 min in a mixture of n-butanol, pyridine, acetic acid, and water (15:10:3:12 v/v).

### Inside-out patch-clamp recordings

Electrophysiological recordings were performed as described (20). Patch pipette solution contained 136 mM NMDG-Cl, 2 mM MgCl_2_, 5 mM HEPES, pH=7.4 with NMDG. Bath solution contained 134 mM NMDG-Cl, 2 mM MgCl_2_, 5 mM HEPES, 0.5 mM EGTA, pH=7.1 with NMDG. MgATP (Merck, A9187) was added from a 400 mM aqueous stock with pH adjusted to 7.1 using NMDG. P-ATP was added from a 10 mM aqueous stock (Biolog LSI, P 012). Purified rPKA-C and bPKA-C were added from 40-60 μM stock solutions; final phosphate concentration following bPKA-C addition was ∼1 mM. VX-770 was added from a 10 μM stock in DMSO. Recordings were done in a continuously flowing bath solution, bath composition could be exchanged with a time constant of ∼20 ms using electronic valves (ALA-VM8, ALA Scientific Instruments). Membrane potential was −40 mV, recording temperature was 25°C. Currents were amplified and filtered at 2 kHz (Axopatch 200B, Molecular Devices), digitized at a sampling rate of 10 kHz (Digidata 1322A, Molecular Devices), and recorded to disk (pCLAMP 9, Molecular Devices).

### Data analysis and statistics

Macroscopic CFTR currents were Gaussian-filtered at 50 Hz and baseline-subtracted (pCLAMP 9, Molecular Devices). Mean steady-state currents under various test conditions were normalized to the mean current observed in the presence of 2 mM ATP + 300 nM bPKA-C in the same patch. Half-times for current activation (t_1/2_) were estimated as the time required for reaching 50% of the final, steady-state current amplitude.

Data are presented as mean±S.E.M., with the number of experiments indicated in each figure legend. Significances were evaluated using Student’s t-test.

---

**Figure S1.**
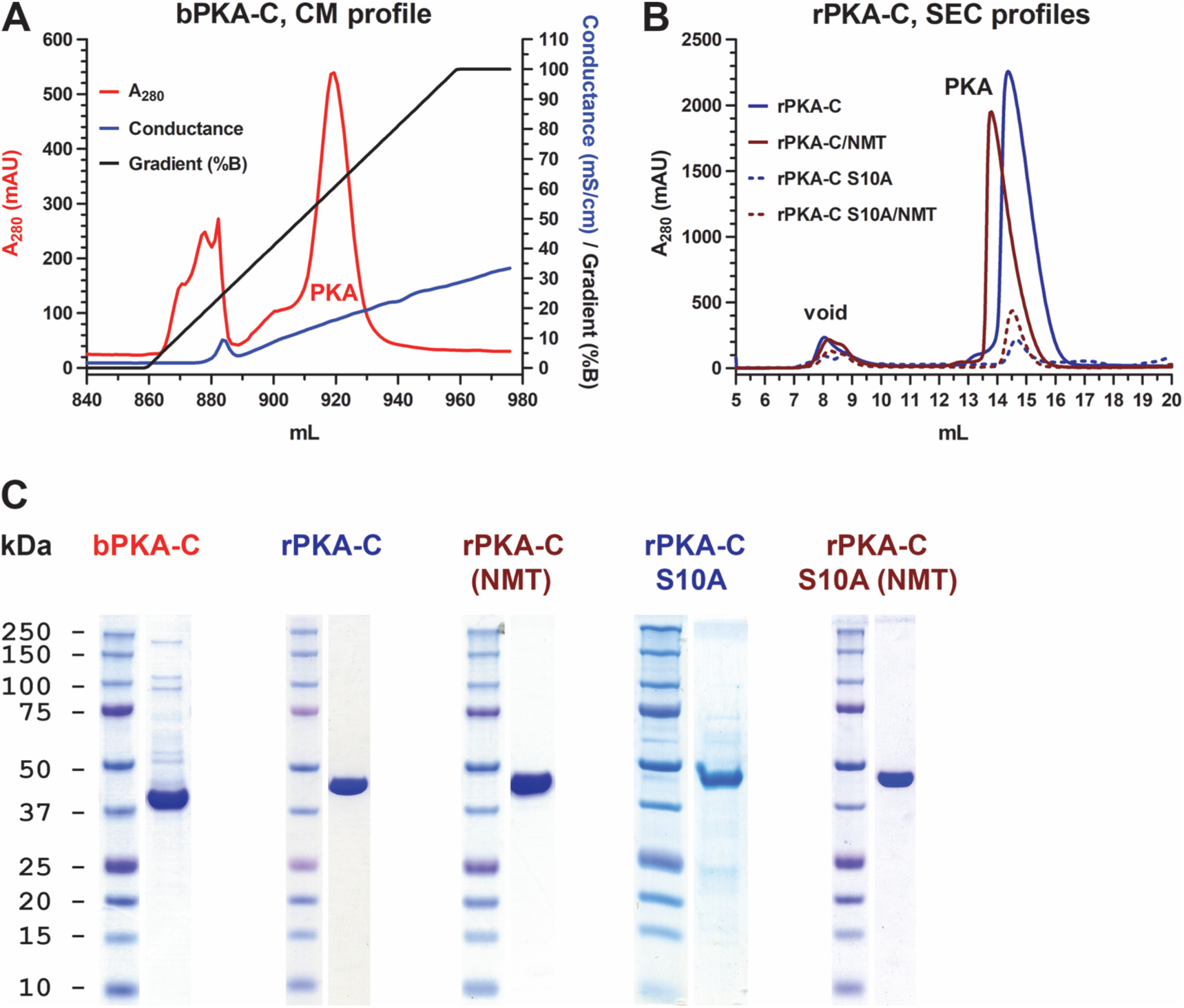
Chromatograms and final purity of PKA-C variants. *A*, Final cation-exchange chromatography (CM Sephadex C-50, Sigma-Aldrich C50120) elution profile, detected as absorbance at 280 nm, of bPKA-C (*red*). Applied salt gradient (*black*; 10 to 300 mM potassium phosphate, pH=6.9) and specific conductance of eluate (*blue*, mS/cm) are plotted. *B*, Size-exclusion chromatography (SEC; Superdex 200 10/300 GL, GE Healthcare Hungary) elution profiles, detected as absorbance at 280 nm, of WT rPKA-C (*dark blue*), WT rPKA-C co-expressed with NMT (*brown*), S10A rPKA-C (*dark blue dashed*), and S10A rPKA-C co-expressed with NMT (*brown dashed*). *C*, Coomassie-stained SDS PAGE gels of bPKA-C, WT rPKA-C, WT rPKA-C co-expressed with NMT, S10A rPKA-C, and S10A rPKA-C co-expressed with NMT. Molecular weights for marker ladder bands (*Precision*, Bio-Rad) are labeled in kDa.

**Figure S2.**
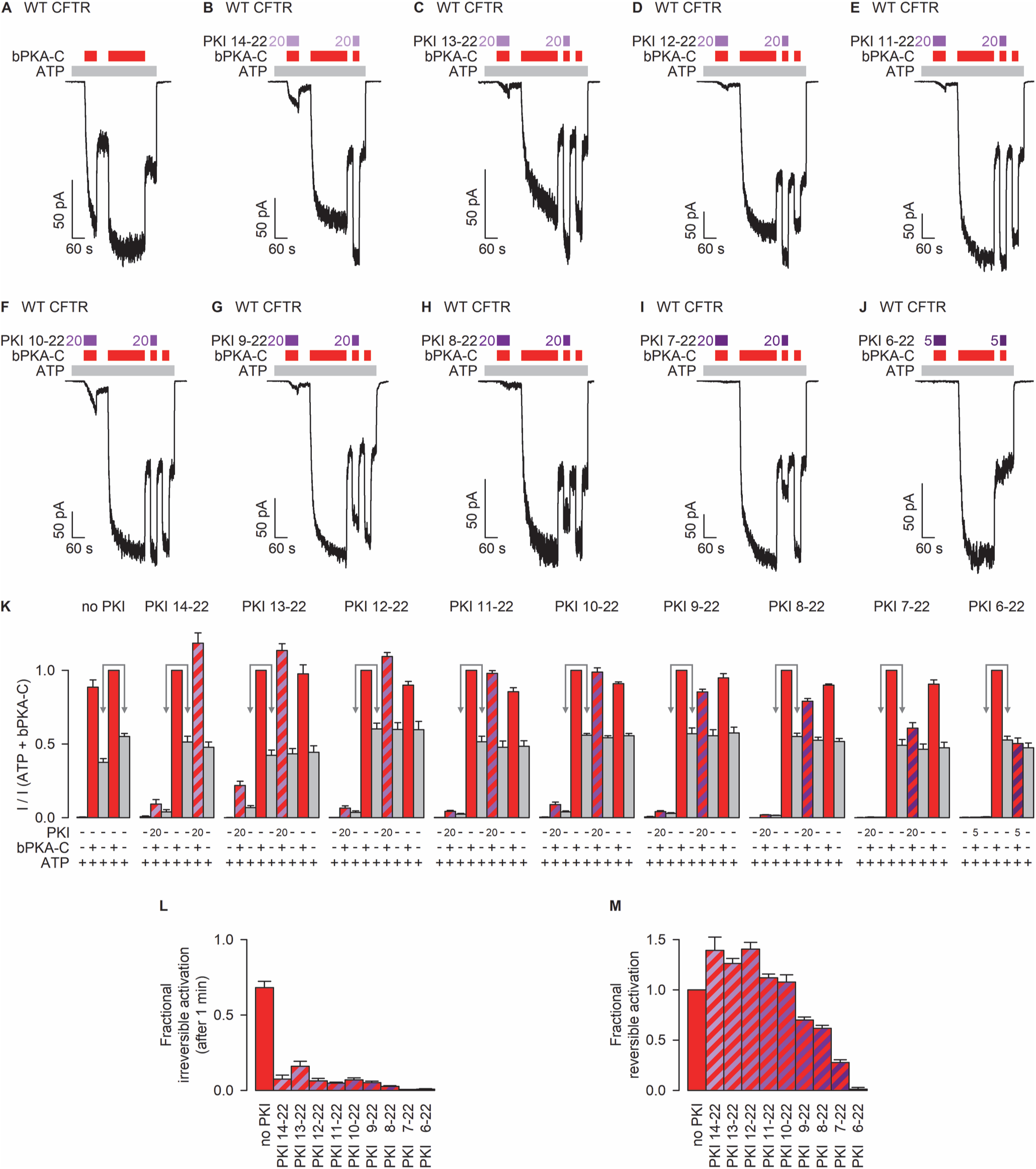
Narrowing down the region on PKI responsible for blocking reversible CFTR activation. *A-J*, WT CFTR currents activated by repeated exposures to 300 nM bPKA-C (*red bars*) in the presence of 2 mM ATP (*light gray bars*), with or without PKI peptides of various lengths (*purple bars*). Colored numbers indicate PKI concentrations in μM. Panels *A*, *B*, *J* replot Figs. 1G, I, H, respectively, for comparison. *K*, Steady-state currents in the sequential segments of the experiment in panels *A*-*J*, normalized to that in the presence of ATP+bPKA-C (*red column*; (*A*, *2nd red column*)). *L*, Fractional CFTR phosphorylation after 1 minute in 300 nM bPKA-C alone or together with PKI peptides of various lengths, reported as the ratio of the two *gray columns* marked by the *double arrows* in panel *K*. *M*, Fractional reversible stimulation by 300 nM bPKA-C in the absence and presence of PKI peptides of various lengths. Bars plot (I(test)-I(ATP))/(I(ATP+bPKA-C)-I(ATP)), where the “test” condition represents exposure to ATP+bPKA-C + PKI. Data in *K*-*M* represent mean±SEM, n=6-11.

**Figure S3.**
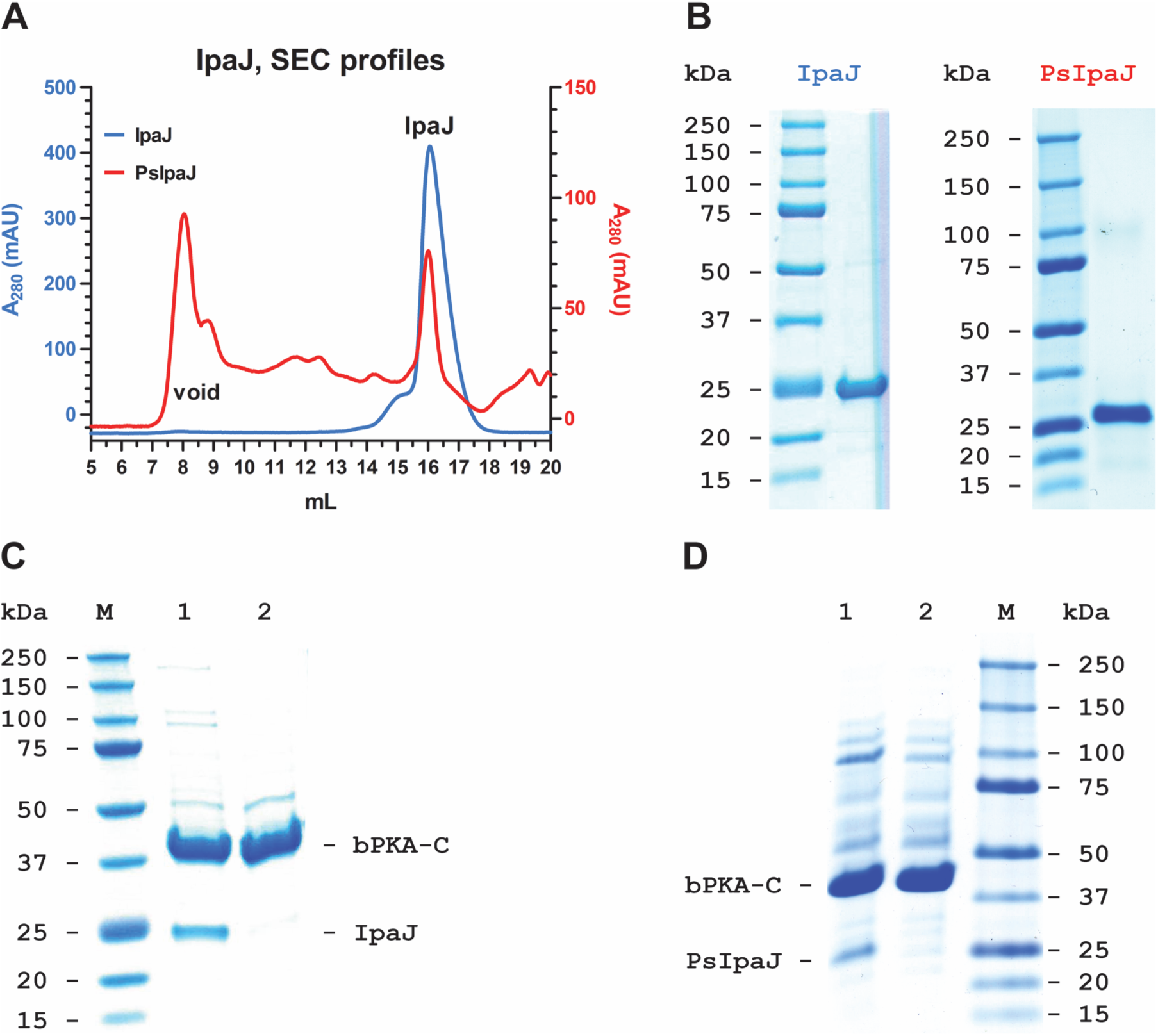
Assessment of purity for IpaJ, PsIpaJ, and IpaJ-treated bPKA-C. *A*, Size-exclusion chromatography (SEC; Superdex 200 10/300 GL, GE Healthcare Hungary) elution profiles, detected as absorbance at 280 nm, of IpaJ (*blue*) and PsIpaJ (*red*). *B*, Coomassie-stained SDS PAGE gels of IpaJ (*left*) and PsIpaJ (*right*). *C*-*D*, Coomassie-stained SDS PAGE gels of the de-myristoylation reaction mix (lanes 1) containing bPKA-C and either IpaJ (*C*) or PsIpaJ (*D*), and of bPKA-C purified from the cysteine protease following treatment (lanes 2). In panels *B*-*D* Molecular weights for marker ladder bands (*Precision*, Bio-Rad) are labeled in kDa.

**Figure S4.**
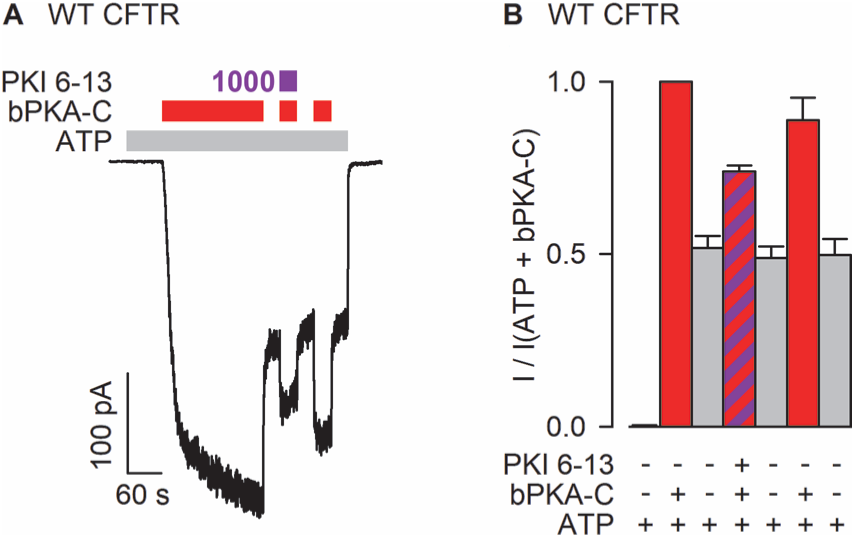
PKI N-terminal helix blocks reversible CFTR current activation by bPKA-C. *A*, Macroscopic current through inside-out patch excised from *Xenopus laevis* oocyte injected with cRNA for WT CFTR. Channels are activated by repeated exposures to 300 nM bPKA-C (*red bars*) in the presence of 2 mM ATP (*light gray bar*), with or without 1000 μM PKI(6–13)amide (*dark purple bar*). Membrane voltage was −40 mV. *B*, Steady-state currents in the sequential segments of the experiment in panel *A*, normalized to that in the presence of ATP+bPKA-C (*1st red column*). Data represent mean±SEM, n=7. Currents in the presence of ATP+bPKA-C + PKI(6–13)amide were significantly smaller (p=7.3·10^-5^) than the average of the currents in the bracketing segments in ATP+bPKA-C.

## Notes

### Competing Interest Statement

The authors have declared no competing interest.

